# Dynamic cognitive representations in the dorsal pallium of adult zebrafish

**DOI:** 10.64898/2026.05.19.726250

**Authors:** Kim Palacios-Flores, Jan Eckhardt, Kuo-Hua Huang, Sriram Narayanan, Rainer W. Friedrich

## Abstract

Brains rely on internal models of the world to interpret sensory input and to simulate the future. In the mammalian hippocampal-entorhinal network, environments are represented by cognitive maps that contain spatially selective neurons such as place, grid, head direction and object-vector cells^1–6^. Neurons with allocentric spatial tuning have recently been discovered also in non-mammalian organisms including larval zebrafish^7^ but it remains unclear to what extent these neurons establish internal cognitive representations. We measured neuronal activity in telencephalic area Dc of head-fixed adult zebrafish exploring a novel, richly structured virtual reality. Neurons were sharply tuned to one or multiple locations and collectively represented environmental space. Activity fields exhibited neuron-specific associations to visual landmarks, indicating a prominent vectorial component in spatial representations. Population activity evolved and became increasingly informative as fish explored the environment. When landmarks were removed after familiarization, landmark-associated activity partially persisted and subsets of neurons reported prediction errors, implying that activity was in part driven by an internal representation. Strong functional coupling among neuronal ensembles and winner-take-all dynamics suggest that representations evolve by refinements of pre-structured networks. The teleost brain therefore generates internal models of structured environments that are optimized by experience and enable cognitive inference and prediction.

## Introduction

Cognition relies on internal models of the world that are established by evolution and by an individual’s experience. Concepts of such “cognitive maps” have been strongly influenced by representations of physical space in hippocampal-entorhinal networks of mammals^8–10^. These networks contain neurons such as place cells^1^, grid cells^2,3^, head direction cells^4^, object-vector cells^5,6^ and other functional neuron types^11–14^ that collectively represent an environment and the location of an animal within it. During navigation, network activity is updated on a moment-to-moment basis by integrating multisensory inputs from internal (e.g., proprioceptive input evoked by body movements) and external sources (e.g., visual input evoked by movements relative to landmarks), tracking the animal’s trajectory over time. Depending on brain state, the same networks can also generate population activity that is not directly driven by sensory input and encodes information about past trajectories^15^, future trajectories^16^ or objects that have been removed from an environment^11^. Hence, hippocampal-entorhinal networks represent relations between an animal and its environment also during memory recall, prediction, or planning, implying retrieval of information from internal models during cognitive brain functions. Moreover, hippocampal-entorhinal networks can establish cognitive maps of non-spatial information^17^ and mediate episodic memory^18,19^. Studies using more complex environments and tasks indicate that neurons frequently exhibit mixed selectivity and conjunctive responses to different types of inputs^20–24^. Spatial cognitive maps in the hippocampal-entorhinal loop have thus become a leading model to study biological neuronal networks underlying memory and cognition.

Computational models show that latent representations of physical space with place cell-like units may, in principle, be generated by different mechanisms and network architectures^25–31^. In biological networks, cognitive maps of spatial environments have been described primarily in mammals. Recently, however, neurons with allocentric spatial tuning – referred to as place cells – were discovered also in the forebrain of food-caching birds^32^. Moreover, representations of boundaries, head direction, and locomotion variables were found in the goldfish telencephalon^33,34^, and an abundance of place cells was found in the telencephalon of larval zebrafish^7^. Zebrafish diverged from the mammalian lineage approximately 450 million years ago, yet place cells in larval zebrafish integrate spatial information from visual cues and self-motion, and their place fields remap in different environments^7^, as observed in mammalian hippocampus. In addition, neurons encoding heading direction, which is critical for updating positional information during navigation, have been found in the anterior hindbrain of zebrafish larvae^35–37^. The larval zebrafish telencephalon therefore records an animal’s position as it navigates an environment using functional neuron types related to those found in mammals. However, further information is needed to examine whether spatial representations in mammals and teleosts share common algorithms and mechanisms. Importantly, it remains to be determined to what extent the teleost brain establishes internal, memory-based representations of environments that can be accessed in the absence of sensory input to support prediction and other cognitive computations.

We studied spatial maps of visual environments in the telencephalon of adult zebrafish using a high-resolution closed-loop virtual reality (VR). We focused on a distinct subregion of pallial area Dc, which is thought to be phylogenetically related to mammalian cortex^38,39^. To determine how representations of environments are shaped by experience we examined neuronal population activity as animals repeatedly explored an initially novel, naturalistic corridor with visual landmarks. We discovered internal representations of environments that evolved with experience. These representations were characterized by a continuum of place fields covering environmental space, by the systematic organization of spatially selective fields as a function of landmarks’ positions in the VR, and by the generation of complex responses upon landmark deletion. The latter comprised landmark-related memory traces and prediction error signals, sometimes conjunctively represented within individual neurons. Highly correlated and anticorrelated activity fluctuations in different subsets of spatially selective neurons point to an underlying mechanism that involves nonlinear mapping of visual and self-motion information by a structured, possibly pre-configured network.

### Spatially selective neurons in cDc

We analyzed representations of spatial environments in the telencephalon of adult zebrafish, which have ∼100 times more neurons than larvae and exhibit a broader repertoire of cognitive behaviors^40^. Head-fixed adult zebrafish were swimming through a linear virtual corridor made of naturalistic textures (rocks, pebbles) with symmetrical, visually complex landmarks (2D images of aquatic creatures) on the walls (Fig. 1a). Four variations of this corridor were used that differed in the number and spatial arrangement of visual landmarks: 7Reg (seven regularly spaced landmarks), 5Reg (five regularly spaced landmarks), 5Irr (five irregularly spaced landmarks), and 0Lm (no landmarks). Navigation was restricted to the forward direction, and fish were teleported back to the beginning of the corridor after each traversal (trial; Supplementary Video 1; 48 ± 31 s/trial [mean ± s.d.]; median: 43 s; inter-quartile range: 57 s – 29 s). Fish had no prior experience with these or similar environments and no rewards or punishments were delivered.

**Fig. 1.**
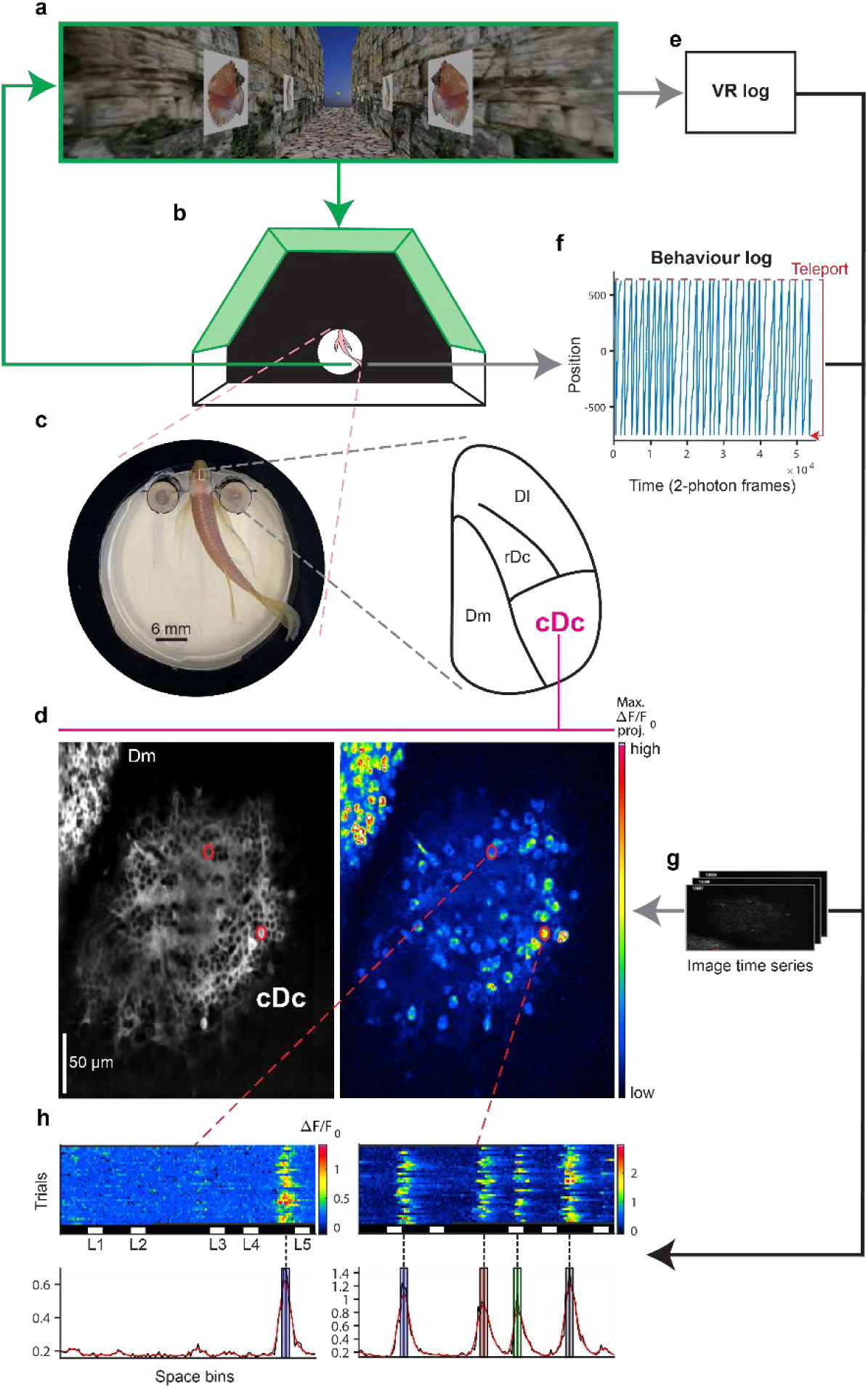
Experimental design for the detection of structured neuronal activity in cDc. **a,** Fish perspective of a VR environment with landmarks. **b,** Schematic of a head-fixed adult zebrafish in a semi-hexagonal tank for VR projection. Light green areas correspond to the walls where the VR is projected. Green arrows represent closed loop VR update based on tail movements. **c,** Head-fixed adult zebrafish. Right: schematic dorsal view of the right telencephalic hemisphere. Dl, lateral zone; Dm, medial zone; rDc, cDc, rostral and caudal regions of the central zone. **d,** Time-averaged raw fluorescence (left) and maximum projection of **Δ**F/F_0_ time series (right) in the same view. Scale bar, 50 μm (left). **e, f, g,** Schematic: data acquisition and analysis workflow. **h,** Top: spatial activity maps of successive trials of two neurons circled in **d**. Blanked bins (dark gray) correspond to periods outside the time of data acquisition (first and last rows) or to periods of mechanical instability that were excluded from analysis. White rectangles below the color map indicate positions of landmarks (L1-L5). Bottom: Trial-averaged neuronal activity. Black: unfiltered; red: low-pass filtered. Black vertical lines depict peaks of activity fields; colored bars depict ±2 space bins around peaks. Same conventions are used in related plots throughout all figures.

Neuronal activity was measured by 2-photon calcium imaging through the intact skull using a transgenic line that expressed the calcium indicator GCaMP6f throughout the dorsal telencephalon (Methods). Measurements were targeted to a distinct subregion in the caudal part of telencephalic area Dc, referred to as cDc, that was identified reliably across individuals based on salient anatomical landmarks^41,42^ (Fig. 1c,d, Methods). The field of view contained, on average, 320 ± 53 somata per fish (mean ± s.d., n = 14 fish) (Fig. 1d). Activity (ΔF/F_0_) was recorded during 15 – 30 min long sessions (Fig. 1g). In a subset of fish (n = 4), activity was recorded in two sessions separated by 40 – 147 min (see below). In two additional fish used for landmark deletion experiments (see below), activity was recorded in 4 or 5 sessions. The closed-loop configuration of the VR was maintained throughout experiments (Fig. 1a,b,f). Each fish was exposed to one virtual environment (7Reg: 2fish; 5Reg: 4 fish; 5Irr: 6 fish, 0Lm: 2 fish). Activity measurements therefore captured early stages in the formation of representations of novel environments.

Subsets of neurons showed pronounced activity at specific locations in the virtual corridor (Fig. 1h). To systematically identify spatially modulated neurons, we spatially binned activity in each trial to obtain *spatial activity maps* and identified peaks that were statistically significant across trials (Methods). The spatial information content of neurons with at least one significant peak was then quantified using a well-established measure of spatial specificity (Methods). Only neurons whose spatial specificity exceeded the expectation based on shuffle controls by at least three standard deviations were further analyzed for statistical significance of activity peaks on a trial-by-trial basis (Methods). We defined *activity fields* as the spatial extent exceeding half-height around each significant peak. Neurons with at least one activity field were defined as spatially selective (Fig. 1h).

Out of 3311 analyzed neurons of fish exposed to VR configurations with landmarks (n = 10 fish), 461 (14%) were spatially selective (Fig. 2a, Extended Data Fig. 1a,b), with either one (246 neurons, 53%) or multiple (215 neurons, 47%) activity fields (Fig. 1h, 2e,f; Extended Data Figs. 1c, 2a). In the 0Lm environment, the fraction of neurons with fields was substantially lower (3%) and fields appeared broader (Fig. 2h; Extended Data Fig. 2b). To assess whether cDc neurons primarily represent spatial location or time, we analyzed activity also as a function of time relative to trial onset. Temporal activity maps were less well aligned across trials than spatial activity maps (Fig. 2f,g) and temporal specificity was significantly lower than spatial specificity of the same neurons (*P* = 2.08 x 10^-73^, Wilcoxon Signed-Rank test) (Fig. 2a, inset). We therefore conclude that cDc neurons represent primarily spatial rather than temporal information.

**Fig. 2.**
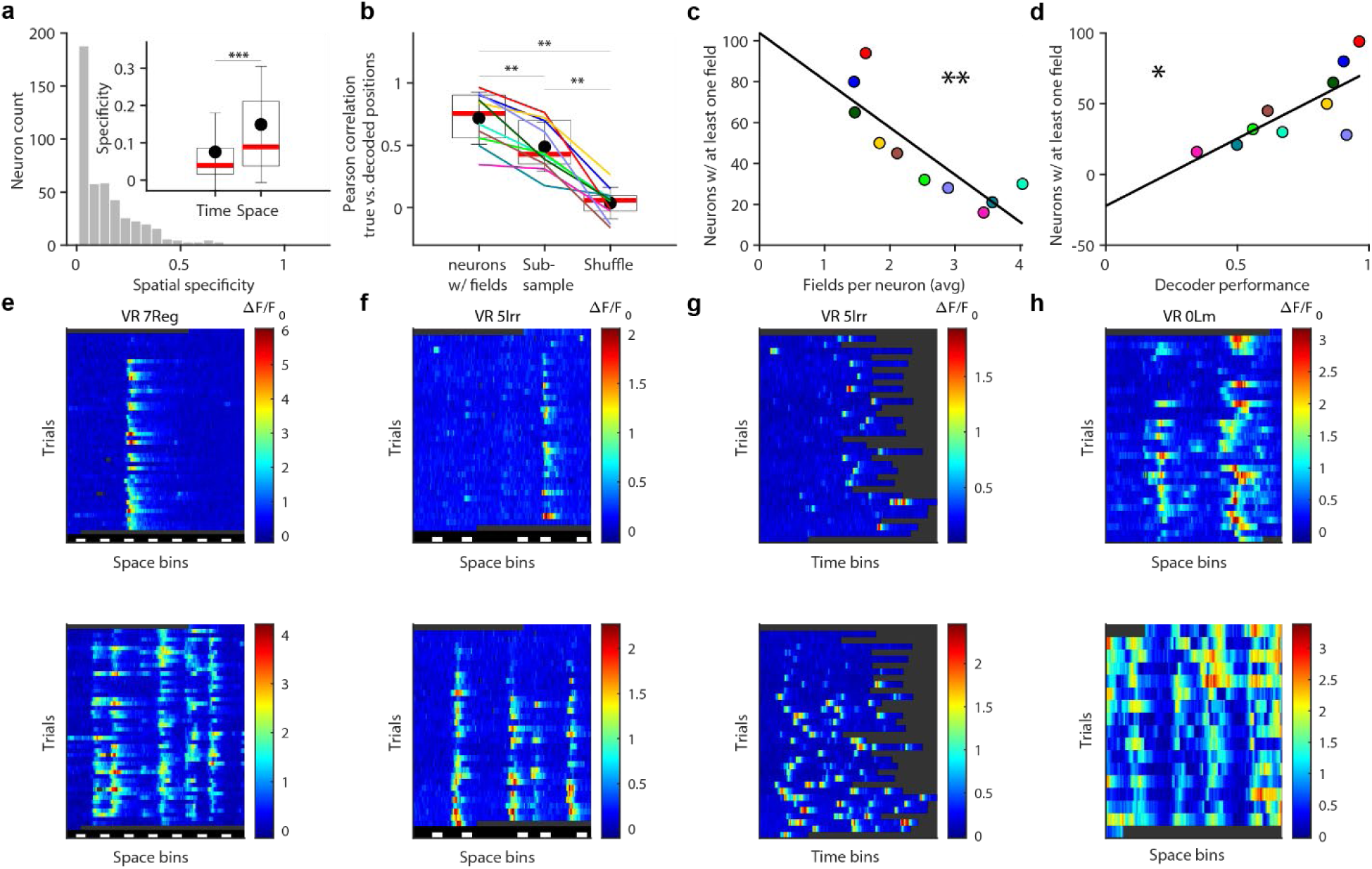
Spatial selectivity of neuronal activity across VR environments. **a,** Histogram of spatial specificity for all neurons with at least one field across all 10 fish exposed to VRs with landmarks. Inset compares specificity of activity binned spatially or temporally. In this and subsequent plots, dots and error bars show mean and s.d., box plots show median and surrounding quartiles. **b,** Pearson correlation between decoded and true fish positions along the VR. Left: decoding based on the activity of all neurons with fields in each fish; center, decoding after subsampling spatially selective neurons to equalize neuron counts across fish; right, decoding based on shuffled activity traces. Colored lines show data from individual fish. **c,** Number of neurons with activity fields as a function of fields per neuron across fish. Color code as in **b**; line shows linear fit. **d,** Number of neurons with activity fields as a function of decoder performance (mean correlation). Color code as in **b** and **c**; line shows linear fit. **e**, **f,** Spatial activity maps of neurons imaged simultaneously in the 7Reg and 5Irr environment, respectively. Top: single-field neurons; bottom: multi-field neurons. **g,** Same neural activity data as in **f** but binned along the temporal dimension. **h,** Spatial activity maps of two neurons from fish in the 0Lm environment. Conventions as in Fig. 1h.

We next trained a neural network to decode spatial position from neural activity on a trial-by-trial basis in environments with landmarks (Methods). For each fish, we used the full set of spatially selective neurons, as well as subsampled sets of neurons (n = 16) to equalize counts across fish. In both cases, decoding performance was significantly higher than expectations derived from shuffled activity traces (Fig. 2b, Extended Data Fig. 1d), confirming that population activity enables positional decoding in individual fish. We further observed that fish with a higher number of spatially selective neurons had, on average, fewer fields per neuron (Pearson correlation coefficient = −0.84, *P* = 0.0024) (Fig. 2c). This translated into a positive correlation between the number of spatially selective neurons and decoder performance (Pearson correlation coefficient = 0.76, *P* = 0.0103) (Fig. 2d) and a negative correlation between decoder performance and the average number of fields per neuron (Pearson correlation coefficient = −0.66, *P* = 0.04) (Extended Data Fig. 1e). These results indicate that cDc networks exhibit predispositions for representations of novel environments with variations between individuals.

### Representations of structured environments

Our observations from VR environments with and without landmarks indicate that landmark-driven inputs contribute significantly, although not exclusively, to the spatial selectivity of cDc neurons, consistent with the contribution of both visual and self-motion information to spatial selectivity in the mammalian hippocampus and neocortex^43,44^. To further examine the influence of landmarks we first analyzed the autocorrelation function of spatial activity maps in two environments with different landmark configurations, 7Reg and 5Irr. If activity is driven by landmarks, the average autocorrelation should exhibit a modulation corresponding to the autocorrelation of the landmark pattern. Consistent with these expectations, autocorrelation functions of spatial activity maps had multiple peaks at locations predicted by the landmark autocorrelations (Fig. 3a,b).

**Fig. 3.**
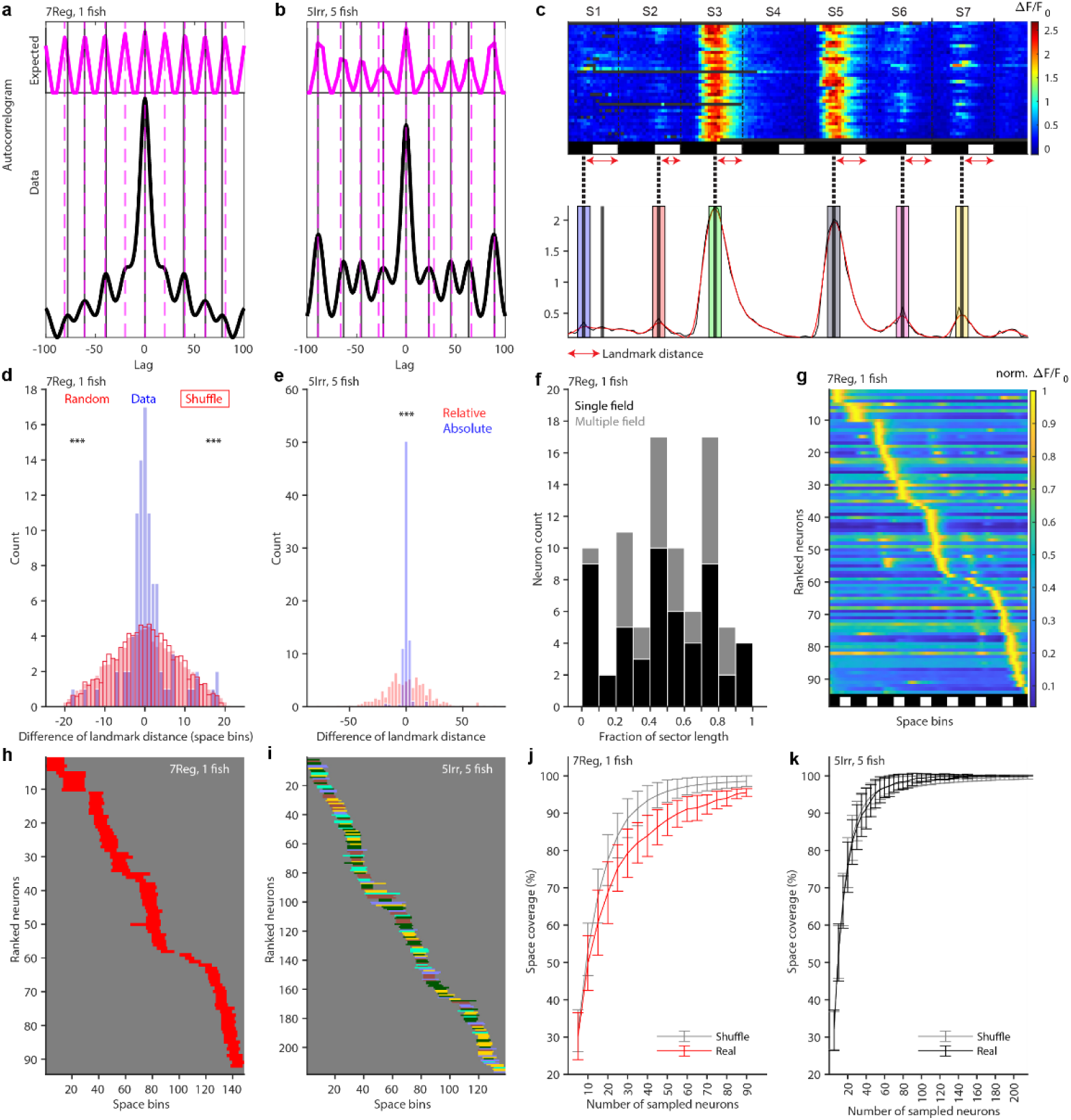
Space coverage at the neural population level across VR environments. **a,** Neural activity autocorrelogram averaged across neurons of one fish behaving in the 7Reg VR (black curve) and expected autocorrelogram based on landmark positions only (magenta curve). Vertical black and magenta lines depict peaks of corresponding autocorrelograms. **b,** Same as **a** for neurons pooled across all 5 fish behaving in the 5Irr VR. **c,** Spatial activity map (top) and field detection (bottom) of one neuron in the 7Reg environment. Conventions as in Fig.1 h. The vertical line without a surrounding colored bar depicts a detected peak that did not pass the statistical significance threshold for field detection. Dashed black lines on the spatial activity map indicate sectors defined by landmarks (S1 – S7). Red arrows indicate landmark distance. Conventions as in Fig. 1h. **d,** Histogram of pairwise differences in landmark distances, calculated for all fields of each multi-field neuron from one fish in the 7Reg environment (blue). Red: shuffled controls (null distributions; filled red bars: randomly drawn distance values; red outline: randomized neuron labels in each sector). **e,** Histograms of pairwise differences of landmark distance values expressed as absolute distances (blue) or as fractions of sector length (red) for all 5 fish behaving in the 5Irr VR. **f,** Histogram of landmark distances, expressed as a fraction of sector length, for single-field (black) and multi-field (gray) neurons from one fish in the 7Reg environment. **g,** Normalized activity of all spatially selective neurons from one fish in the 7Reg environment, normalized per neuron and sorted by peak location. **h,** Binarized fields extracted from data in **g**. **i,** Binarized fields of neurons from 5 fish in the 5Irr environment, one color per fish as in Fig. 2b. **j,** Space coverage by binarized fields from one fish in the 7Reg environment as a function of neuron number (red) and shuffle control (gray). Error bars show s.d.. **k,** Same as in j for all 5 fish in the 5Irr environment (black) and shuffle control (gray).

The autocorrelation indicates that activity fields of individual neurons are linked to landmarks, but it does not reveal the precise spatial relationship between fields and landmarks. To address this question, we divided each corridor into non-overlapping sectors that were delimited by the end of each landmark. Each activity field was then assigned a “landmark distance” defined as the distance between the activity peak and the end of the corresponding sector (Fig. 3c). The majority of neurons had at most one activity field per sector. In some cases, two fields were observed in the same sector, in particular in the 5Irr environment in sector 3, where the distance between landmarks was large. In this environment, the maximum landmark distance analyzed was limited to the length of the smallest sector. For each neuron with multiple fields, we subtracted landmark distances of different fields from each other (Methods), resulting in a narrow distribution of differences that was centered on zero by design (Fig. 3d). This experimentally observed difference distribution was compared to two null distributions. The first null distribution was constructed by computing differences between randomly drawn distance values within the permitted sector length. The null distribution was significantly broader than the experimentally observed distribution (*P* =1.19 x 10^-5^, two-sample Kolmogorov-Smirnov test). These results imply that landmark distances were consistent within neurons, in accordance with results of the autocorrelation analysis. The second null distribution was constructed as described after shuffling of neuron labels across fields within corresponding sectors. This null distribution was also significantly broader than the experimentally observed difference distribution (*P* = 3.92 x 10^-5^, two-sample Kolmogorov-Smirnov test) (Fig. 3d), demonstrating that landmark distances differ between neurons.

To further examine whether activity fields represent absolute distances or the relative position (“phase”) between landmarks we computed difference distributions using both metrics in the 5Irr environment where landmarks are spaced irregularly. Difference distributions were significantly narrower when distances were quantified using the absolute metric (*P* = 1.27×10^-31^, two-sample Kolmogorov-Smirnov test) (Fig. 3e), indicating that activity fields of cDc neurons represent characteristic absolute distances to landmarks. Similar distance-dependent firing fields have been observed in the medial entorhinal cortex of mice navigating a linear VR^13^ in neurons that are likely to correspond to object-vector cells in 2-dimensional environments^5^. Our results therefore indicate that vectorial information contributes strongly to representations of spatial environments in cDc.

To examine how activity fields covered space within sectors we analyzed the distribution of characteristic landmark distances across neurons and sectors. The distribution was broad without an obvious pattern (Fig. 3f), indicating that cDc neurons collectively cover a wide range of landmark distances, spanning the entire sector space. To examine spatial coverage of the full environment we sorted spatial activity maps by their maximum amplitude field and found that activity fields across the neural population covered the entire corridor (Fig. 3g). To further characterize spatial coverage, we generated a binarized representation of space by thresholding each neuron’s highest-amplitude field at half maximum (Fig. 3h and i). Hence, each spatially selective neuron contributed one segment with a defined location and width to the binarized representation of the corridor. We then randomly subsampled neurons and analyzed space coverage as a function of neuron number (Methods). The increase in space coverage with neuron number was not changed substantially after randomizing the positions of binary segments (Fig. 3j,k). Hence, the distribution of activity fields along the corridor was almost indistinguishable from a random distribution, indicating that activity fields collectively covered the environment without obvious patterning by landmarks or other discontinuities.

### Evolution of spatial representations

Activity fields of cDc neurons changed in intensity and, in some cases, appeared or disappeared within individual imaging sessions, implying that the representation of the virtual environment evolved over time. To analyze this evolution over more extended periods of time we measured activity from the same neurons during two imaging sessions separated by 32 – 174 trials (40 - 147 min; n = 4 fish; fish A, 7Reg, 174 trials; fish B, 5Reg, 81 trials; fish C and D, 5Irr, 54 and 32 trials, respectively). Between sessions, fish continued to swim through the VR but no activity data were recorded. Activity fields of each neuron were identified independently in sessions 1 and 2 and assigned a common identifier (field ID) when they overlapped in space (Methods). 265 neurons had at least one field in at least one of the sessions. In the majority of these neurons (n = 222; 84%) the number of fields changed between imaging sessions (Fig. 4a). 72 neurons (27%) had fields only in session 1 while 99 neurons (37%) had fields only in session 2 (Fig. 4a). From a total of 500 fields, 180 fields (36%) were lost after session 1, 170 fields (34%) appeared de novo in session 2, and 150 fields (30%) were detected in both sessions. The number of neurons with one or two fields increased while the number of neurons with 3 or more fields decreased (Fig. 4b). Consequently, the fraction of single-field neurons increased (Fig. 4c) while the mean number of fields per neuron decreased (*P* = 0.0133, Wilcoxon rank sum test) (Fig. 4d), even among neurons with fields in both sessions (*P* = 1.88 x 10^-4^, Wilcoxon Signed-Rank test) (Fig. 4e). Examples of activity profiles evolving with time are provided in Fig. 4f–l and Extended Data Figure 3a. In some cases, we were able to capture the emergence of activity fields from a background devoid of spatial selectivity within one continuous imaging session. Examples of these transformations occurring as abrupt transitions are presented in Fig. 4k and l and Extended Data Fig. 3 (top row, fourth spatial activity map, S1). These observations underscore the dynamic nature of the internal representation.

**Fig. 4.**
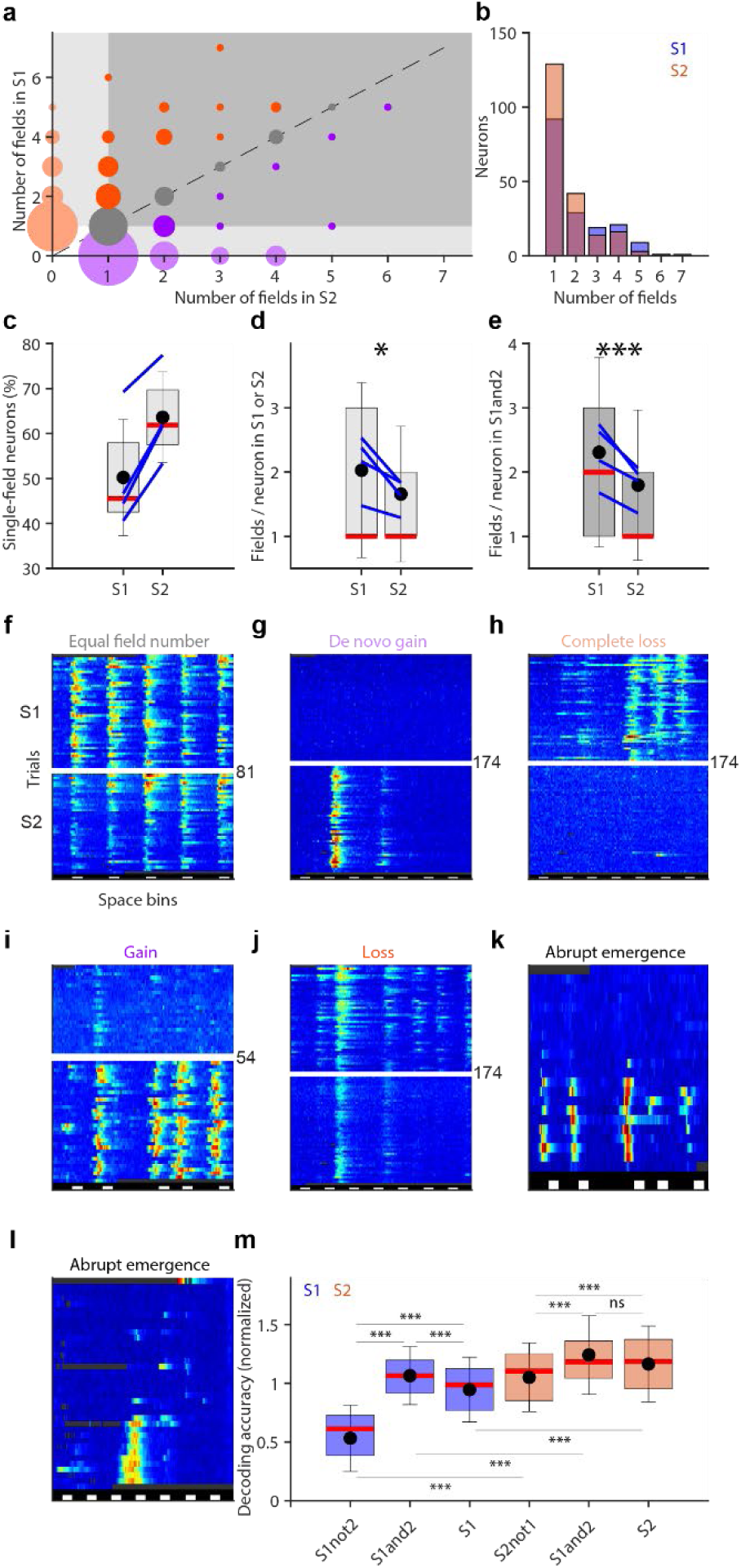
Dynamics of the internal representation. **a,** Number of fields per neuron in sessions 1 and 2 (S1, S2; circle area is proportional to neuron number; n = 4 fish). Colors highlight complete loss of fields (orange), partial loss of field number (red), and gain of fields (purple; pale: *de novo* gain of fields). Light gray area covers neurons with fields in S1 *or* S2, dark gray area covers neurons with fields in S1 *and* S2. **b,** Histogram showing the absolute number of neurons with different numbers of fields in S1 and S2. **c,** Percentage of single-field neurons in S1 and S2. Lines connect individual fish. **d**,**e,** Number of fields per neuron in the subset of neurons with fields in S1 *or* S2 (**d**) and S1 *and* S2 (**e**). Lines connect means of individual fish. **f** - **j,** Spatial activity maps of individual neurons in S1 and S2 illustrating different types of changes. S1 and S2 are separated by a white gap; number of trials between sessions is indicated. Conventions as in Fig. 1h. **k**, **l,** Spatial activity maps showing the abrupt emergence of spatial selectivity within an imaging session in different environments. **m,** Normalized decoding accuracy in S1 and S2 using different subsets of spatially selective neurons (100 instances of the decoding network, each trained on a different subset). To account for differences in neuron numbers across fish, decoding accuracy (Pearson correlation between decoded and real positions) was normalized to the mean decoding accuracy across all subsets in each fish (n = 3; one fish was excluded because neuron number in one category, S2not1, was too low). In **c**, **d**, **e**, and **m**, dots and error bars show mean and s.d., respectively; box plots show median and surrounding quartiles.

To determine how the evolution of population activity affected decoding of spatial information we trained artificial neural networks to decode position based on different sets of spatially selective neurons (Methods): neurons that lost their fields after session 1 (S_1NOT2_), neurons that had fields only in session 2 (S_2NOT1_), neurons with fields in both sessions (S_1AND2_), and all neurons with fields in one of the two sessions (S1: union of S_1NOT2_ and S_1AND2_; S2: union of S_2NOT1_ and S_1AND2_). Neuron numbers were equalized across sets by subsampling to the largest possible number (size of the smallest set; minimum of 8 neurons) in each fish (n = 3 fish; 14, 14, and 8 neurons, respectively) and decoding accuracy was quantified by the Pearson correlation between decoded and true positions (Fig. 4m; Extended Data Fig. 3b–d). Overall, decoding accuracy increased significantly from session 1 to session 2. Accuracy was lowest based on neurons that lost their fields after session 1 (S_1NOT2_), significantly higher based on neurons that gained fields in session 2 (S_2NOT1_), and highest based on neurons with fields in both sessions (S_1AND2_). The reorganization of population representations therefore enhanced positional information on timescales of minutes to hours, consistent with an increasingly more uniform coverage of a physical environment by place cells observed in freely swimming zebrafish larvae^7^. The observed dynamics of activity patterns supports the hypothesis that the brain establishes a cognitive map of the environment that is optimized by experience.

### Cognitive inference and error signals in cDc

We next asked whether cDc generates internal representations of environments that enable cognitive inference and prediction. To address this question, we examined whether spatial representations organized as a function of landmarks can remain accessible in the absence of landmark-driven sensory input. We obtained neural activity data of the same neurons over multiple imaging sessions (Fig. 5a–e). In the first session, fish were familiarized with the virtual environment for 20 min (∼20 trials). During subsequent sessions we deleted all landmarks for five consecutive trials after trial 8 (n = 7 deletions in 2 fish). We refer to each deletion/neuron pair as a “deletion event”. For each deletion event we compared activity immediately prior to (8 trials) and during deletion of landmarks (5 trials). As a control, we analyzed size-matched blocks of consecutive trials from experiments without landmark deletions (“mock deletions”, 7 fish, one mock deletion per fish). In a small subset of deletion events, landmark removal resulted in intense activity that was typically more broadly distributed than activity fields related to landmarks. Deletion events showing this type of activity were identified by thresholding of the mean activity and analyzed separately (see below). Among the remaining deletion events, landmark deletion abolished spatially modulated activity in the majority of cases. In a subset of cases, however, we detected activity during landmark deletion that was spatially aligned with locations corresponding to activity fields in the presence of landmarks, indicating that landmark-related activity partially persisted.

**Fig. 5.**
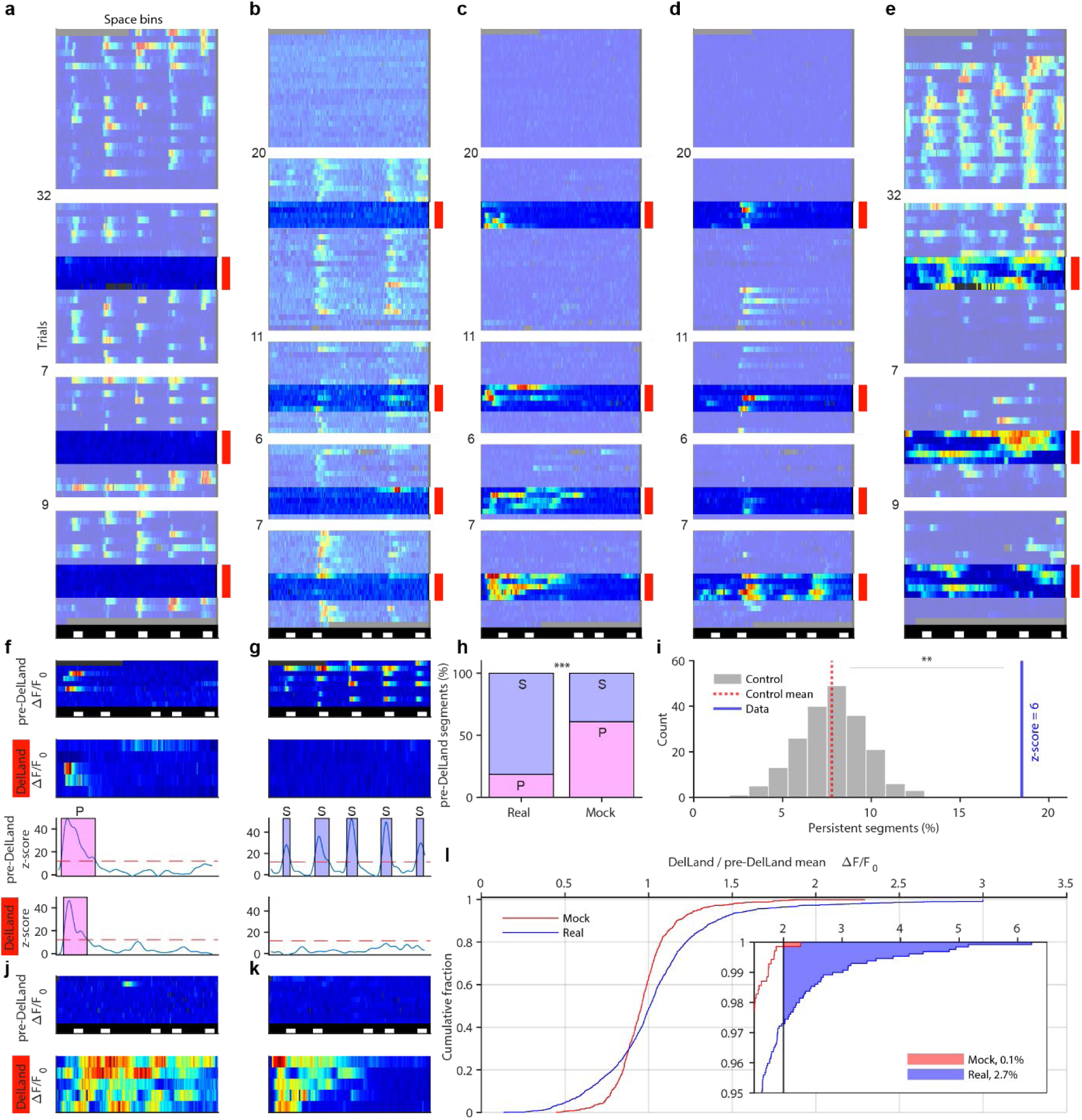
Consequences of landmark deletion on neuronal activity. **a** - **e**, Spatial activity maps of individual neurons across multiple imaging sessions including landmark deletions (DelLand). Sessions are separated by white spacing, numbers indicate trials between sessions. Red bars and stronger colors highlight trials when landmarks were deleted. Conventions as in Fig. 1h. **f**, **g,** Top: spatial activity maps of trials before and during landmark deletion containing activity segments classified as persistent (P; **f**) or sensitive (S; **g**). Conventions as in Fig. 1h. Bottom: mean z-scored activity. Shaded areas depict activity segments; dashed line shows threshold for detection. **h,** Percentage of sensitive and persistent activity segments in real and mock deletions, pooled across all deletions and fish. **i,** Distribution of the percentage of persistent activity segments observed in shuffle controls (gray, red dashed line shows mean) compared to the observed percentage of persistent activity segments (blue line, z-score indicated). **j, k**, Spatial activity maps before and during landmark deletion of two neurons showing activity classified as error signals. **l,** Cumulative distribution of activity ratios (mean activity during/before landmark deletion) and corresponding distribution for mock deletions. Distribution is clipped at ratios ≥ 3 in the main graph; inset shows enlargement of the full upper range. Black vertical line depicts threshold used to identify error signals. Percentage of error signals among all deletion events is given by the areas under the curves (colored).

To quantify the persistence of landmark-related activity we defined activity fields using a slightly different procedure (mean activity over trials significantly exceeding baseline in ≥ 5 consecutive spatial bins; Methods) to account for the low number of trials before and during landmark deletion. We refer to spatially defined significant activity detected using this procedure as *activity segments*. An activity segment detected prior to landmark deletion was classified as “persistent” (Fig. 5f) when it overlapped with an activity segment during landmark deletion and “sensitive” otherwise (Fig. 5g). The probability of observing persistent activity segments by chance was quantified by the same procedure after circularly permuting activity in trials without landmarks (“spatial randomization”, Methods).

Eighteen percent of activity segments (n = 160/866 from 643 analyzed neurons in 2 fish) were classified as persistent after landmark deletion whereas 61% of activity segments (n = 523/859 from 2438 analyzed neurons in 7 fish, *P* = 1.69×10^-72^, Chi-square test of independence) persisted in the presence of landmarks (mock deletions) (Fig. 5h). Hence, landmark deletion confirmed that landmark-driven sensory input contributes significantly to spatially selective activity in cDc. Moreover, spatial randomization significantly decreased the fraction of activity segments classified as persistent from 18% to 8% (mean ± s.d. over 200 randomizations; *P* = 0.005) (Fig. 5i). Activity during landmark deletion therefore contained traces of prior activity segments and, thus, information about landmark-related activity. These results indicate that the activity of cDc neurons contains spatial information that is inferred from an internal, memory-based model of the environment.

We next examined high-intensity responses to landmark deletion in more detail. To detect overall changes in activity during landmark deletion (Fig. 5j and k), we normalized the mean activity during each deletion event to the mean activity during the 8 trials prior to landmark deletion. As expected, decreases in activity were frequent and more prominent than in mock deletions because a significant subset of activity segments were sensitive (Fig. 5l). In a subset of deletion events, however, landmark deletion increased activity beyond the largest increases observed in mock deletions (Fig. 5l). We identified such high-intensity responses using a threshold of twice the mean activity before landmark removal, corresponding approximately to the largest activity change observed in mock deletions. Activity exceeded this threshold in 2.7 % of deletion events (35/1294) during landmark deletion but only in 0.1 % of mock deletions (1/697) (Fig. 5l), implying that high-intensity activity was triggered by the removal of landmarks. We therefore interpret high-intensity responses as error signals encoding a mismatch between the predicted and the experienced appearance of the environment. Since the computation of a prediction requires an internal model, the observation of error signals further supports the conclusion that cDc neurons retrieve information from internal representations of the environment.

Error-related activity was often broad but not uniformly distributed throughout the corridor (Fig. 5c,d,e). Error signals were observed in neurons that were largely inactive prior to landmark deletion (Fig. 5c,d), but also in neurons showing spatially selective activity (Fig. 5e). Hence, at least some cDc neurons may conjunctively encode information about prediction error and location. Additional activity profiles illustrating the variety of responses elicited upon landmark deletion are shown in Extended Data Fig. 4. In summary, landmark deletions revealed inference of landmark-based spatial relationships and prediction error computations in cDc. As these computations require access to an internal representation of the environment, our observations imply that cDc performs memory-based cognitive inference.

### Functional neuronal ensembles in cDc

Spatial selectivity may emerge in a broad spectrum of networks with different architectures including generic, initially unstructured recurrent networks and networks with specific pre-configured connectivity^25–31,45^. In highly structured networks, subsets of neurons may be expected to form functional ensembles because they receive similar feed-forward and/or recurrent inputs. To explore this possibility we examined correlated variability of neuronal activity. Spatially selective activity of many cDc neurons varied substantially across trials, sometimes switching abruptly between high activity and silence in successive trials (Fig. 6f–n). To examine whether these variations were coordinated across the population we quantified pairwise trial-to-trial (noise) correlations between the signal magnitude of all fields in each of the 10 fish exposed to VRs with landmarks (96 ± 28 fields per fish, mean ± s.d.). As a control, we performed the same analysis after shuffling trial order independently for each field. Shuffling significantly reduced the probability of observing large positive or large negative noise correlations and resulted in a narrower overall distribution of correlation coefficients (*P* = 1.51 x 10^-290^, two-sample Kolmogorov-Smirnov test; n = 100 shuffles; Fig. 6a), demonstrating that high trial-to-trial activity correlations occurred more frequently than expected by chance.

**Fig. 6.**
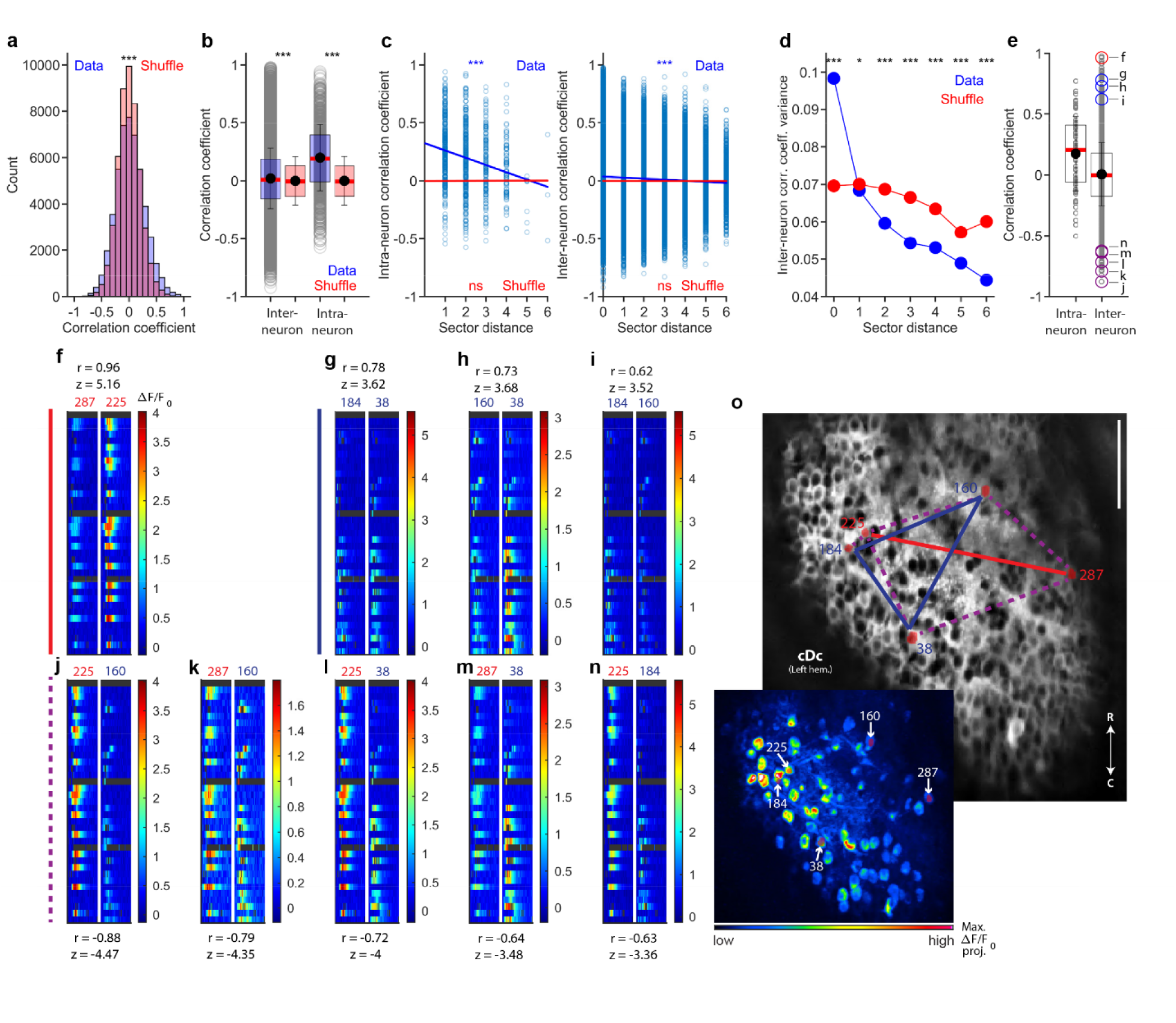
Coordinated “noise” correlations in the neural population. **a,** Histograms of pairwise trial-by-trial (noise) correlations between all possible field pairs (n = 10 fish in environments with landmarks; blue) and shuffle controls (red). **b,** Inter- and intra-neuron noise correlations (blue) and shuffle controls (red). **c,** Intra- (left) and inter-neuron (right) noise correlations as a function of sector distance. Blue and red lines show linear regressions to data and shuffle controls, respectively. **d,** Variance of inter-neuron noise correlations as a function of sector distance. **e,** Intra- and inter-neuron noise correlations in one fish in the 7Reg environment. Colored circles depict high positive and negative correlations corresponding to the examples shown in **f** – **n**. **f** – **n**, Subregions of spatial activity maps centered on fields from different pairs of neurons with high positive or negative noise correlations highlighted in **e**. Conventions as in Fig. 1h. IDs of neurons are given by numbers above each activity map. Correlation coefficients and z-scores are indicated above (positive correlations) or below (negative correlations) each plot. **o,** Raw fluorescence image (time-averaged) and regions of interest (ROIs) corresponding to highlighted neurons. Solid lines depict high positive correlations, dashed purple lines depict high negative correlations; different colors represent different ensembles. Left hem., left hemisphere; R, rostral; C, caudal; scale bar, 50 μm. Inset shows maximum projection of the **Δ**F/F_0_ time series in the same field of view and depicts locations of ROIs. In **b** and **e**, dots and error bars show mean and s.d., respectively; box plots show median and surrounding quartiles.

We next compared the covariation of activity between different fields of the same neurons (intra-neuron correlations; 1008 field pairs) and between fields of different neurons (inter-neuron correlations; 48’406 field pairs). Inter- and intra-neuron correlations both spanned a broad range and were, on average, significantly higher than correlations between shuffled controls (*P* =1.55 x 10^-51^ and *P* =1.64 x 10^-118^, respectively, Wilcoxon rank sum test) (Fig. 6b). On average, intra-neuron correlations were higher than inter-neuron noise correlations. However, highest positive and negative correlation coefficients were found between fields of different neurons (Fig. 6b).

To examine how noise correlations depend on the spatial separation of fields we assigned each field to its corresponding sector in the corridor and expressed inter-field distances as the difference in sector number (“sector distance”). Intra-neuron noise correlations decreased significantly with sector distance (Pearson correlation = −0.22, *P* < 10^-16^), approaching zero at large distances (Fig. 6c, left). In shuffled controls, mean correlations remained near zero for all sector distances. Hence, trial-by-trial covariation of activity in the same neurons was most pronounced between nearby fields. Inter-neuron noise correlations also decreased significantly with sector distance (Pearson correlation = −0.05, *P* < 10^-16^; Fig. 6c, right). Moreover, extreme inter-neuron noise correlations occurred primarily between fields of the same or nearby sectors (Fig. 6c, right). To quantify this observation, we measured the variance of correlation coefficients, which was highest among same-sector correlation coefficients and decreased as a function of sector distance. Compared to shuffle controls, the observed variance was significantly higher in the same sector (*P* = 3.21×10^-136^, two-sample F-test) and significantly lower across sectors (sector distance = 1: *P* < 0.05; sector distances > 1: *P* < 0.001; two-sample F-test; Fig. 6d). These results indicate that subsets of cDc neurons with spatially related activity fields are organized into functional ensembles.

To systematically locate functionally related sets of neurons we searched for core groups of 3 neurons with same-sector noise correlations ≥ 0.6 and added further neurons that were linked to at least one core group member through high absolute noise correlations (|r| ≥ 0.6) (Methods). As illustrated by the example in Fig. 6e–o, groups consisted of distinct ensembles of neurons that were defined by their correlation structure (e.g., neurons [287, 225] and [38, 160, 184]). Within ensembles, spatially selective activity co-fluctuated across trials (Fig. 6f–i) whereas across ensembles, trial-to-trial variations were anti-correlated, resulting in almost mutually exclusive activity on different trials (Fig. 6j–n). Additional examples of functional ensembles are shown in Extended Data Fig. 5. In all cases, functionally coupled neurons were distributed throughout cDc without an obvious topography. Spatially selective population activity in cDc is therefore organized, at least in part, into distributed functional ensembles that are activated in a winner-take-all fashion. These observations are consistent with inhibitory interactions between neuronal ensembles^46^ and possibly excitatory coupling within ensembles, suggesting highly structured connectivity. It appears unlikely – albeit not impossible – that such connectivity emerged *de novo* from a random network during the earliest explorations of the environment, when activity could not be systematically analyzed. We therefore hypothesize that sensory and self-motion inputs are initially integrated by pre-configured networks and that these networks are then refined by experience-dependent plasticity to establish increasingly more specific internal representations.

## Discussion

Our results revealed cognitive spatial maps of structured environments in an anatomically well-defined subregion of telencephalic area Dc in adult zebrafish. cDc neurons had one or multiple sharply delineated activity fields that may be referred to as place fields and collectively tracked an individual’s position. Spatially selective activity was strongly reduced but not abolished in the absence of landmarks, indicating that positional information is computed primarily from relations to landmarks with some contribution from path integration. Consistent with this conclusion, individual neurons represented specific distances to landmarks. In mice, similar distance coding in linear virtual environments is characteristic for neurons such as object-vector cells in medial entorhinal cortex^5,13^. In two-dimensional environments, these and related neuron types represent distances and directions towards objects, giving rise to a vectorial code of space that is thought to be critical for navigation^6,12,14,47^. Our results thus provide evidence for vectorial representations of spatial environments in teleosts, although further experiments are required to examine spatial selectivity of cDc neurons in two-dimensional environments.

Previous experiments in zebrafish revealed neurons with place fields in the telencephalon of freely swimming^7^ but not head-fixed zebrafish larvae. Potentially important distinctions of our approach from previous studies using head-fixation include (1) the use of adult zebrafish, (2) the use of a high-resolution VR with naturalistic textures and objects, and (3) an efficient algorithm for closed-loop VR updates that supports naturalistic swimming^42^.

Under these conditions, spatially selective activity was reliably observed in Dc, which has been loosely linked to isocortex^38,39^. Functional properties of cDc neurons are reminiscent of cue-locking, object-vector and object-trace cells in the medial and lateral entorhinal cortex^5,11,13^, suggesting that cDc may be related to entorhinal cortex. However, cDc neurons also share functional features with neurons in other brain areas including landmark vector and place cells in hippocampus^1,12,14,43^. Further studies are therefore needed to explore phylogenetic and functional relationships between cDc and mammalian brain areas.

Few previous studies examined the emergence of spatial cognitive maps during the initial exploration of a novel environment. Consistent with observations in rodents^48–50^ we found prominent spatially selective activity covering the entire environment already at early stages of exploration. Subsequently, representations were reorganized on a timescale of minutes to hours, resulting in increased selectivity of individual neurons and more informative population activity. While the spatial position of activity fields remained stable, their signal magnitude changed over time. Individual fields could appear de novo, as observed in hippocampal neurons after current injection^51^, or disappear abruptly between trials. These observations are consistent with models of pre-configured circuits that mediate initial spatial representations and a subsequent refinement of functional connectivity by experience, possibly by a self-organizing process that optimizes coding efficiency^45,52,53^.

We found small functional ensembles of co-tuned cDc neurons that were identified by a striking covariability of activity fluctuations. Instances of highly correlated activity within ensembles, but anti-correlated across ensembles, indicate that population activity exhibited winner-take-all dynamics consistent with cross-ensemble inhibition^46^, further supporting the notion of pre-configured circuit structure. In theory, such an organization is consistent with computational models that explicitly assume inhibition between small and distinct computational modules^30^. The small size of the zebrafish brain offers the opportunity to address structural predictions of such models by combining activity measurements with connectomics^54^.

Manipulations of the virtual environment allowed us to directly examine whether the zebrafish brain establishes internal representations of an environment that enable computations underlying cognitive processes such as inference and prediction. Consistent with this hypothesis, we found traces of previous spatially selective activity after landmark deletion, implying that landmark-based spatial relationships were inferred from an internal model. Moreover, landmark deletion triggered prominent error signals, which requires the comparison of sensory input to an internally generated expectation^55–57^. Furthermore, conjunctive responses of individual neurons are consistent with the notion that internal representation and prediction error computations are closely inter-related, as one requires the other. Taken together, our results show that the zebrafish brain generates internal models of structured environments that are accessed for cognitive computations, consistent with a conservation of higher brain functions and the underlying mechanisms across vertebrates.

## Methods

### Animal models

Experiments were performed in adult (6 – 18 months old) Tg(neuroD:GCaMP6f) zebrafish^58^ (Danio rerio) of either sex in a nacre background. Animals were raised and kept under standard laboratory conditions (26 – 27 °C, 13 h/11 h light/dark cycle). All animal procedures were approved by the Veterinary Department of the Canton Basel-Stadt, Switzerland.

### Head-fixation

Head fixation was performed as described in Huang et al.^42^. Briefly, adult zebrafish were anesthetized by immersion in tricain methanesulfonate (MS-222; 0.03%) and held in a custom-build gyroscopic device. Anesthesia was maintained throughout the surgical procedure by perfusion with fish water containing MS-222 (0.01%) through a cannula inserted into the mouth. After removing the skin over the telencephalon and adjacent bones, light-weight, stainless-steel head bars made from syringe needles were attached to specific bones of the skull using tissue glue and dental cement. Fish were mounted by gluing the pins to a pair of vertical posts integrated into the floor of a transparent semi-hexagonal tank for submersion in the VR. After mounting, the anesthetic was washed out and fish recovered from anesthesia within ∼1 min.

### Virtual reality

VR experiments were performed as described^42^ with some modifications (see below). The tank with the mounted fish was placed under a custom-built multiphoton microscope with resonant scanners controlled by custom-written software based on Scanimage^58,59^. Infrared LEDs were placed outside the tank to acquire images of the tail with a video camera from below at a resolution of 90 x 70 pixels (medio-lateral x rostro-caudal) and a frame rate of 50 Hz. The VR scene was projected onto the three planar walls of the fish tank using three projectors.

The shape of the tail was tracked continuously (50 Hz) using custom LabVIEW code based on Huang et al.^42^. Forward movement was inferred from image frames based on the curvature and undulation of the caudal part of the tail as described^42^. Using custom software written in Python, the inferred forward movements were used to update the projection of the VR scene and to deploys user-defined protocols such as transient landmark deletions (see below).

The Python-based game engine Panda3D was used to update the location of the virtual cameras in the VR. A set of three virtual cameras was implemented to capture three adjacent 45° x 60° (height by width) fields of view side by side (total of 45° x 180°, height by width). The geometry of the virtual world was constructed using the 3D modelling software Blender.

Some modifications were made compared to the VR codebase described previously in Huang et al.^42^. The basic control of virtual cameras as a function of behaviour (tail images) was implemented as described^42^ but the underlying Python codebase was redesigned for modular, combinatorial, and scalable control of all aspects of the VR geometry. This was achieved through the implementation of a Finite State Machine architecture which deploys VR protocols within an experimental data acquisition and trial structure framework (https://github.com/fmi-basel/DynCogRep-VR). Additionally, the new codebase incorporates functionality for recording all relevant VR events that occur during an experiment, generating a detailed log for subsequent data analysis.

After initiating the closed loop VR, the first ∼15 minutes were used to fine-adjust the gain of forward movement for each individual fish as in Huang et al.^42^ and to locate the region of interest in the dorsal telencephalon (see below). The closed-loop VR was maintained throughout this period and the remainder of the experiment.

All VRs were variations of a linear corridor with naturalistic textures and images on the walls. Fish movement was restricted to the forward direction within the corridor without allowing for turns.

### Calcium imaging

Calcium imaging was performed using a modified 2-photon microscope (Sutter MOM scope) equipped with a x16 Nikon objective as described^42^. Images were acquired using 8 kHz resonant scanners at a frame rate of 30 Hz. To prevent contamination of the calcium imaging data by photons from the VR, the LEDs of the projectors and the fluorescence-detecting PMT were temporally gated to alternate between illumination and data acquisition during successive lines (line frequency, 8 kHz), yielding an effective resolution of 512 x 256 pixels. GCaMP6f fluorescence was imaged through the intact skull using an excitation wavelength of 920 nm, a 510/50 nm band-pass emission filter, and 35 mW - 57 mW of power at the sample.

Activity data were acquired in imaging sessions of 15 – 30 min. Data consisted of either a single imaging session per fish, two imaging sessions per fish (evolution of responses from S1 to S2 experiments), or 4 - 5 imaging sessions per fish (landmark deletion experiments). Between imaging sessions, the closed-loop VR was maintained but no fluorescence data were acquired.

Frames from each continuous two-photon imaging session were registered using cross-correlation based on the discrete Fourier transform and adaptive templating. Occasionally, small consecutive sets of frames could not be aligned accurately due to small, transient frame shifts, presumably caused by mechanical perturbations, e.g., during strong tail beats. Frames affected by such events were detected based on the image shifts during alignment and excluded from analysis (blanked). Thresholds to detect transient image shifts for blanking were adjusted to not detect slow image drifts^42^ that could be corrected by alignment. Corresponding activity measurements were blanked in displays of activity data. Data from continuous imaging sessions were then divided into trials corresponding to corridor traversals.

### Location of cDc

Calcium imaging data were acquired from an optical plane containing 320 ± 53 somata (mean ± s.d.) in the caudal subregion of the central zone of the dorsal telencephalon (cDc, as identified in Huang et al.^42^). This distinct region was identified reliably across animals based on two anatomical landmarks: (1) the sulcus ypsilonformis, which delineates the border between the medial zone of the dorsal telencephalon (Dm) and other telencephalic brain areas including cDc, and (2) an obvious anatomical boundary visible in the neuroD:GCaMP6f expression pattern that separates cDc from the rostral subregion of Dc and the lateral zone of the dorsal telencephalon (Dl). This boundary appears to correspond to a boundary in parvalbumin expression^41^. The optical plane was located between 10 μm and 60 μm below the dorsal surface of cDc (mean, 34 μm) in either hemisphere.

### Prediction and verification of neuronal regions of interest (ROIs)

To identify regions of interest (ROIs) corresponding to individual somata, we first generated anatomy maps by computing the pixel-wise mean over aligned frames (excluding blanked frames) of imaging sessions from cDc of different fish. Additionally, we computed maximum **Δ**F/F_0_ projection and local cross-correlation maps. Next, we generated a set of ground truth data by manually delineating cell bodies using such maps. The ground truth data set was then used to train a STARDIST^60^ model. For ROI prediction in STARDIST, we set the prob_thresh and nms_thresh parameters to 0.7 and 0.2, respectively. ROIs predicted by the model were manually verified and adjusted if necessary, and small overlaps between ROIs were programmatically eliminated.

### Mapping of corresponding neurons across imaging sessions

For neural data analyses spanning more than one calcium imaging session we mapped ROIs from one imaging session to data from all other imaging sessions of the same anatomical plane. This mapping included an adjustment of ROI shapes based on elastic alignments of the mean raw fluorescence images from different imaging sessions, which was performed using bUnwarpJ^61,62^. Transformed ROIs were manually verified and adjusted if necessary.

### Activity traces, spatial activity traces, and spatial activity maps

Measurements of fluorescence intensity as a function of time, averaged over all pixels of each ROI, were converted to **Δ**F / F_0_ time traces. F_0_ was defined as the average of the 25% lowest percentile of the fluorescence trace. We refer to these traces as *activity traces*.

Activity traces of each imaging session were averaged in spatial bins along the corridor (10 VR units/bin) to generate *spatial activity traces*. Activity traces were also separated into trials (corridor traversals) prior to spatial binning to visualize activity on a trial-by-trial basis. We refer to these trial-separated, spatially binned matrices as *spatial activity maps*.

In addition, trial-separated activity traces were binned in time windows of 0.33s (10 frames), discarding the first trial in each imaging session because it was usually incomplete. Trial-separated, temporally binned matrices are referred to as *temporal activity maps*.

Some analysis procedures depended on the detection of peaks in spatial activity traces, which may be complicated by noisy fluctuations. To reduce such fluctuations, spatial activity traces were smoothed in some analyses using a low-pass filter (2^nd^ order Butterworth filter; cutoff frequency: 0.25 x Nyquist frequency) and procedures that did not induce phase shifts (MATLAB’s filtfilt command).

For some statistical analyses, **Δ**F/F_0_ measurements were transformed into standardized activity values by subtracting the mean of the 20% lowest **Δ**F/F_0_ values and dividing by their standard deviation. Standardized values therefore represent activity in units of standard deviations (z-scores) relative to an estimate of baseline activity.

### Spatial information

We used the spatial activity traces of neurons with at least one spatial activity field (see below) to quantify spatial information using a well-established measure^63^:

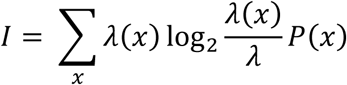

where *I* is the neuron’s spatial information, x is the spatial bin, *P*(x) is the probability that the fish is in spatial bin x, *λ*(*x*) is the mean **Δ**F/F_0_ value when the fish is in spatial bin x, and *λ* is the mean activity of the neuron calculated as *λ* = ∑*_x_ λ*(*x*)*P*(*x*).

The spatial specificity of a neuron has been defined as the spatial information normalized by the neuron’s mean activity^63^:

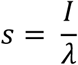

To determine whether a neuron’s activity conveyed significant spatial information we generated a null distribution of spatial specificity values based on 100 shuffled controls. Each shuffled control was generated by circularly permuting activity traces independently in each trial prior to generating the spatial activity trace. A neuron was considered for further analysis (see below) when its spatial specificity exceeded the mean of the null distribution by at least three standard deviations.

### Spatial activity fields

To define spatial activity fields of cDc neurons it is important to take into account the following observations: (1) individual neurons often have multiple fields, (2) trial-to-trial variability of activity can be high, (3) activity within fields may change in intensity during an imaging session, (4) individual fields may appear de novo or disappear during a session. Since commonly used procedures may not robustly detect activity fields under these conditions, we used a multi-step procedure which includes some aspects of previously used methods^7,63^.

Spatial activity traces of each neuron were low-pass filtered as described above, generating a smoothed trace used for detecting all peaks and valleys. The magnitude of each peak was quantified by averaging values within a neighborhood of two space bins in each direction. Peak magnitudes were then standardized relative to the distribution of values in the entire spatial activity map, as described above. Magnitudes therefore represented the activity associated with each peak relative to an estimate of baseline activity in units of standard deviations. Peaks with a magnitude ≥5 were considered as candidates for fields and subjected to further analysis. Neurons without candidate fields were excluded from further analyses.

We then quantified the spatial information of neurons with at least one candidate field. As described above, neurons whose spatial specificity did not exceed the mean of the corresponding null distribution by at least three standard deviations were discarded from any further analyses. In the next step, we quantified the amplitude and width of each candidate field. Amplitude was defined as the value of the smoothed spatial activity trace at the peak after subtracting the mean value of all valleys. Field width was measured as the width at half height around a peak. The spatial extent of a field was defined as the section of the corridor where the smoothed spatial activity trace around a peak exceeded half height, which could be asymmetric. When spatial extents of different candidate fields overlapped, only the candidate field with the maximum amplitude was kept. Fields were required to have positive amplitudes and half-heights within the location of the corridor. The latter was not always the case when fields were located close to the beginning or end of the corridor. Candidate fields that did not fulfill these criteria were excluded from further analysis.

Statistical significance of candidate fields was further assessed using low-pass zero-phase filtered spatial activity map values of each trial. For each candidate field, we searched for peaks within the candidate field’s spatial extent on a trial-by-trial basis. Thus, for each trial, we determined whether the candidate field contained a peak. If so, we quantified its standardized magnitude relative to the distribution of spatial activity map values of the corresponding trial, as well as its amplitude, as described. For each candidate field, we then determined the proportion of trials that fulfilled the following criteria: (1) the spatial extent of the candidate field contained a peak with a magnitude ≥ 7, and (2) the peak had a positive amplitude and decayed to half height on both sides within the corridor. Candidate fields that met these criteria in ≥ 20% of trials were considered spatial activity fields; other candidate fields were discarded from further analyses. Neurons with at least one spatial activity field were considered spatially selective neurons. Visual inspection of data indicates that this procedure selects fields conservatively, i.e., priority is given to minimizing false-positives at the expense of incurring false-negatives.

### Identification of activity fields across imaging sessions

For each neuron, all activity fields detected in session S1 were assigned a unique field ID. Fields detected independently at S2 inherited the ID of a S1 field if their peaks were within a small spatial distance (4 space bins). If necessary, the spatial extent of the field was adjusted to reflect the leftmost and rightmost positions in the union of S1 and S2 fields. When a S2 field overlapped with more than one S1 field it was associated with the S1 field of greatest overlap. If an S2 field had no overlap with any S1 field it was assigned a new field ID. Fields sharing the same ID at S1 and S2 were assumed to be the same field that persisted from S1 to S2.

### Decoding of fish position from neural activity

Decoding was performed using a 4-layer feedforward artificial neural network (ANN) set up and trained using PyTorch. The ANN consisted of an input layer with number of nodes equal to the number of spatially tuned neurons, two hidden layers with 64 and 24 nodes, respectively, and a single output node. Activity traces and fish position from 60% of trials were used to train the neural network and the performance was evaluated on the unseen data. Training was performed over 200 epochs, within which performance was tracked using a subset of held-out trials (10%). This configuration of the neural network was determined empirically; minor changes to the architecture had only minor effects on performance and the overall trends were maintained.

Decoding was performed either by including all spatially tuned neurons or by randomly subsampling neurons to equalize their numbers across fish to match the fish with the lowest count (16 neurons). Frames in which the fish were stationary (velocity <0.01 VR units/s) were excluded from decoding. Performance of the decoder was assessed by computing the average correlation coefficient between the decoded and true positions for a trial and by the average absolute error in decoded and true positions, expressed in units of VR space bins. The performance of the decoder was compared against decoders set up using shuffled activity traces, where the traces were independently shuffled in time for each neuron over a window spanning the length of the recording.

To track how the neural representation of position evolved over time we performed the decoding analysis for experiments in which two recording sessions (S1 and S2) were performed in the same fish. Spatial tuning was determined independently for S1 and S2 and the set of tuned cells was split into the following categories: a) spatially tuned in S1 but not S2, b) tuned in both S1 and S2 and c) tuned in S2 but not S1. The decoding analysis was performed as described above. To make meaningful comparisons between subsets with different neuron numbers, random subsampling was performed to match neuron numbers to the smallest set. Subsampling and decoding were repeated 100 times for each comparison. When comparing the decoding performance of each subset with the pooled population of neurons, the pooled population was subsampled 100 times, keeping the proportion of subsampled neurons the same between each subset in the subsampled set.

### Spatial relationships between activity fields and landmarks

For VRs with landmarks, we divided the VR corridor into non-overlapping sectors. The first sector encompassed the space between the beginning of the corridor and the end of the first landmark. The last sector covered the space between the end of the last landmark and the end of the corridor. The remaining sectors extended from the end of one landmark to the end of the next landmark. The last sector was excluded, as it is bounded by an element different in kind (the end of the corridor).

Each activity field was assigned to a sector based on the location of its peak. The landmark distance of each field was measured as the distance between the peak and the end of the sector. When landmarks were spaced irregularly, landmark distance was limited to a maximum given by the size of the smallest sector. Fields with peaks outside this limit were excluded from analysis. Spatial relationships between fields and landmarks were therefore analyzed using a metric of absolute distance (in VR units) rather than relative distance between landmarks (phase). This metric was chosen because absolute landmark distances were more consistent across multiple fields of the same neurons than phase relationships.

To analyze landmark distances across fields within multi-field neurons we computed the difference between landmark distances for all pairwise combinations of activity fields per neuron. For each subtraction operation, the decision to subtract the smaller from the bigger distance, or vice versa, was performed at random. We then pooled the results across neurons to generate the double-difference distribution.

Two double-difference null distributions were generated for comparison. The first null distribution was generated after replacing the measured landmark distance of each field with a random value between zero and the maximum sector length. The second null distribution was generated after randomly reassigning neuron IDs across fields within corresponding sectors. In both cases, null distributions were based on 100 independent shuffles.

Given the observed consistency of landmarks distances from fields of the same neuron we assigned a characteristic landmark distance to each neuron by computing the mean of the landmark distances of all of its fields.

### Spatial coverage

Spatial activity traces of neurons with at least one field were low-pass filtered as described above, normalized to their maxima, and ranked according to the location of their maxima. For analysis of coverage, we considered only the field with the highest amplitude of each neuron. Spatial activity traces were binarized by setting activity values within the spatial extent of the highest-amplitude field to 1 and all other activity values to 0. Space coverage was then calculated as the percentage of space bins that contained at least one non-zero entry across the population of neurons. Coverage as a function of neuron number was determined by drawing 100 random subsets of neurons for each sample size. As a control, coverage was computed after assigning random spatial locations to each binarized field. This procedure was repeated 100 times for each sample size.

### Landmark deletion

To detect persistent neuronal activity after landmark deletion we compared activity during deletion (5 trials, DelLand window) to activity immediately preceding deletion (8 trials, pre-DelLand window). The analysis was restricted to a small number of trials to minimize the potential influence of activity-dependent refinement processes independent of landmark deletion. Analysis procedures were adapted to conservatively detect activity fields given low trial numbers.

For each deletion event, we computed the spatial activity traces prior to and during deletion by spatial binning of activity traces corresponding to the pre-DelLand and DelLand windows, respectively. Pre-DelLand and DelLand spatial activity traces were then low-pass zero-phase filtered, further smoothed using a running average with a sliding window of five spatial bins, and standardized as described above. We then independently searched the pre-DelLand and the DelLand traces for segments of at least 5 contiguous space bins where standardized activity was ≥12.

Statistical significance of segments was further assessed using low-pass zero-phase filtered spatial activity map values of each trial in the corresponding pre-DelLand or DelLand window. For each segment, we searched for peaks within the segments’ spatial extent on a trial-by-trial basis. Thus, for each trial, we determined whether the segment contained a peak. If so, we quantified its magnitude by averaging values within a neighborhood of two space bins in each direction and standardized it relative to the distribution of spatial activity map values of the corresponding trial, as described. Only segments containing a peak with a standardized magnitude ≥7 within their spatial extent in at least one individual trial were considered for further analysis. We refer to these segments as *activity segments* instead of activity fields because their quantitative characterization is based on a low number of trials.

An activity segment detected in the pre-DelLand window was classified as “persistent” if an overlapping activity segment was detected in the DelLand window. If this requirement was not fulfilled, activity segments were classified as “sensitive” to landmark deletion. Visual inspection of data indicates that this procedure is conservative, i.e., false-positive detections of persistent activity segments appeared to be rare.

To assess whether the fraction of persistent activity segments was statistically different from chance we generated a null distribution based on 100 shuffled controls. Each control was generated by circular permutation of the original activity traces corresponding to each DelLand window trial. Trial-wise circular permutation of DelLand windows was performed for deletion events where the pre-DelLand window contained at least one activity segment.

For some deletion events, we observed that activity during landmark deletion was not spatially localized, as typically observed in the presence of landmarks, but broad and intense (high-intensity activity). To detect such responses, we identified deletion events with at least one activity segment before or during landmark deletion and quantified the ratio of the mean activity trace values during and before deletion. Activity was classified as “high-intensity” in the absence of landmarks relative to the presence of landmarks if the ratio was ≥2. Such responses were considered to be prediction error signals and were analyzed separately.

### Identification of neuronal ensembles

To locate examples of functionally related sets of neurons, we first identified all inter-neuron field pairs complying with two criteria: 1) their trial-wise noise correlation exceeded an absolute threshold (r ≥ 0.6), and 2) fields within each pair belonged to the same VR corridor sector. We refer to these field pairs as “high-pass field pairs”. We then searched for neuron sets of size 3 where each neuron was related to the other two neurons via high-pass field pairs. By identifying and merging trios with at least one shared neuron, we generated a list of unique non-overlapping sets of neurons (“super trios”). Finally, super trios were expanded by adding neurons directly related to any of their members via high-pass field pairs. Such extended super trios correspond to identified functional ensembles.

To locate neurons anticorrelated with functional ensembles, we first identified all inter-neuron field pairs complying with two criteria: 1) their trial-wise noise correlation was ≤ - 0.6, and 2) fields within each pair belonged to the same VR corridor sector. We refer to these field pairs as “low-pass field pairs”. For each ensemble, we identified neurons related to any of its members via low-pass field pairs. Neurons anticorrelated to one ensemble sometimes were strongly correlated among them, forming themselves an ensemble. Additionally, we searched for high-pass field pairs between neurons anticorrelated to the same ensembles to detect associations that did not amount to trio structures.

The above procedure yields sets of neurons related to each other via strong positive or negative correlations. Neurons identified as part of these structures were then subjected to a step of manual curation to exclude potential artifacts such as optical crosstalk. This step involved direct manual inspection of identified neurons and their corresponding ROIs in anatomy and activity maps (maximum **Δ**F/F_0_ projection and local cross-correlation) calculated from image time series. Neurons with potential ambiguities were identified. Next, both high-pass and low-pass field pairs including such neurons were eliminated. These curated lists of field pairs were used as input to the entire neuronal ensemble detection pipeline described above, yielding manually verified functional ensembles.

### Data analysis

Neuronal activity cross-referenced with logged behaviour and VR data was analyzed using custom software written in MATLAB (https://github.com/fmi-basel/DynCogRep-2PVR).

## Author contributions

K.P.F. conceived the project, designed and performed all experiments, developed software, analyzed the data, and wrote the manuscript. J.E. developed software, hardware and provided critical technical support. K.H.H. developed the original methodology and provided critical technical input. S.N. analyzed data. R.W.F. conceived and supervised the project and wrote the manuscript.

## Acknowledgements

We thank E. Arn, P. Argast, and P. Buchmann for technical support, A. Bharioke for helpful discussions, A. Lüthi for comments on the manuscript, and the Friedrich lab members for insightful discussions. This work was supported by the Novartis Research Foundation, by a Novartis Institutes for Biomedical Research Presidential Postdoctoral Fellowship (K.H.H.), by the European Research Council (ERC) under the European Union’s Horizon 2020 research and innovation program (grant agreements no. 742576, 101167289), by the Swiss National Science Foundation (SNSF; grant number 310030_212236) and an EMBO fellowship (S.N.; ALTF 1060-2024).

---

**Extended Data Fig. 1.**
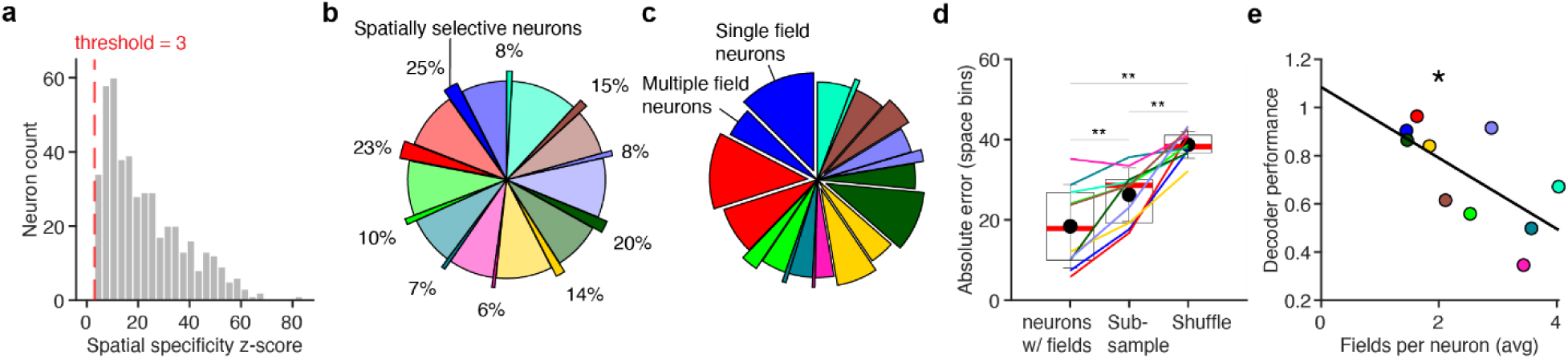
Spatial selectivity of neuronal activity in cDc. **a**, Histogram of spatial specificity z-scores for all spatially selective neurons. The red dashed line represents the imposed significance threshold (z-score ≥ 3) relative to shuffle controls. **b,** Pie chart showing the proportion of neurons contributed by each fish (color coded as in Fig. 2b) to the total of neurons analyzed across all fish. For each fish, the exploded slice (darker shade) and the indicated percentage represent the proportion of neurons with at least one activity field (spatially selective neurons). **c,** Pie chart showing the proportion of neurons contributed by each fish (color coded as in Fig. 2b) to the total of neurons with activity fields across fish. For each fish, the proportion of single field neurons (exploded slice), and of multiple field neurons is indicated. **d,** Absolute error in space bins between decoded and true fish positions along the VR. Left, decoding performed using all neurons with fields for each fish; center, decoding after subsampling spatially selective neurons to equalize neuron counts across fish; right, decoding of shuffled neuronal activity traces. Colors as in Fig. 2b. Dots and error bars show mean and s.d., respectively; box plots show median and surrounding quartiles. **e,** Decoder performance as a function of average number of fields per neuron; line shows linear fit. Colors as in Fig. 2b.

**Extended Data Fig. 2.**
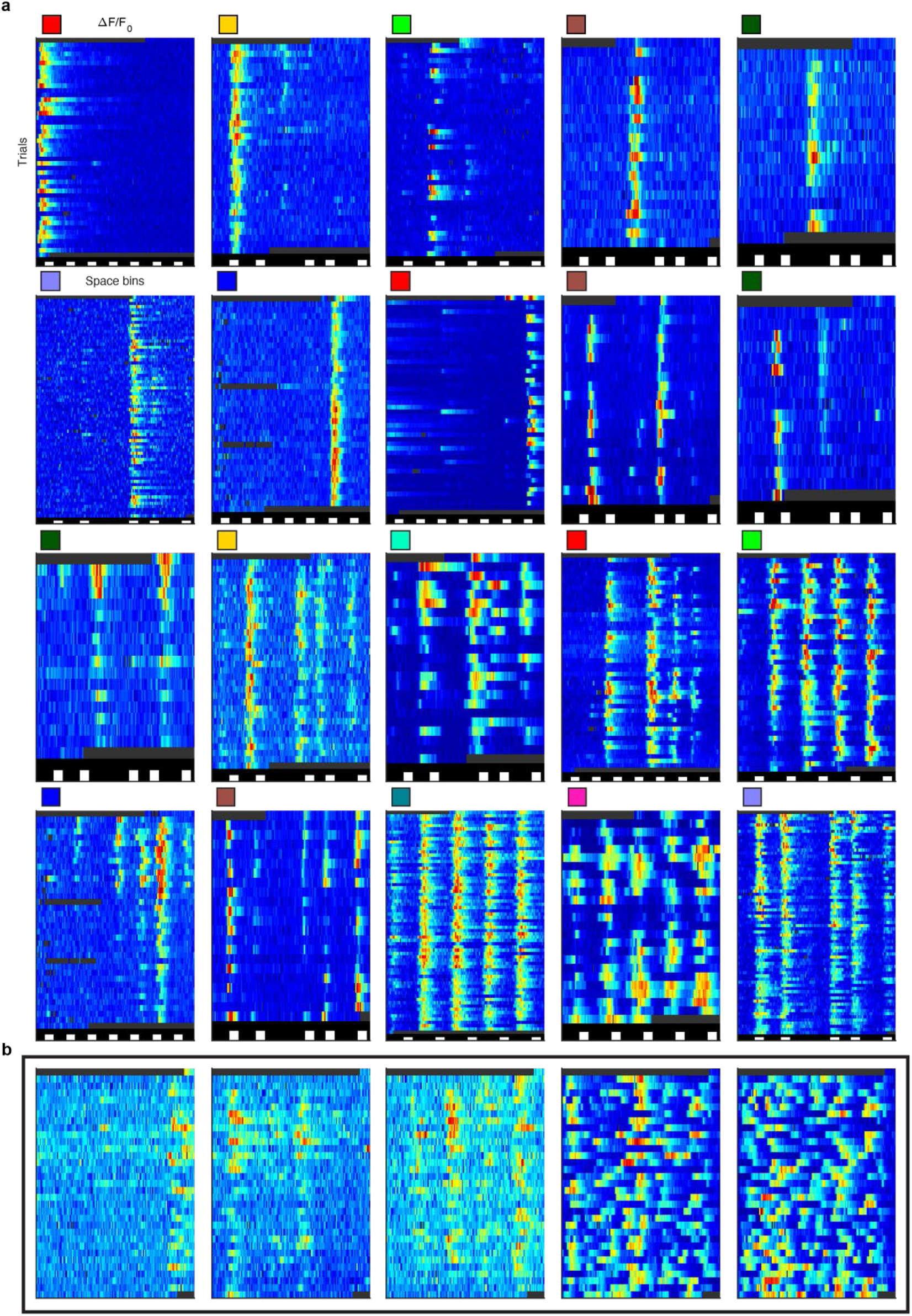
Additional examples of spatially selective neurons. **a,** Spatial activity maps of individual neurons from 10 fish (squares color coded as in Fig. 2b) behaving in different environments containing landmarks. **b,** Spatial activity maps of individual neurons in the 0Lm environment (no landmarks).

**Extended Data Fig. 3.**
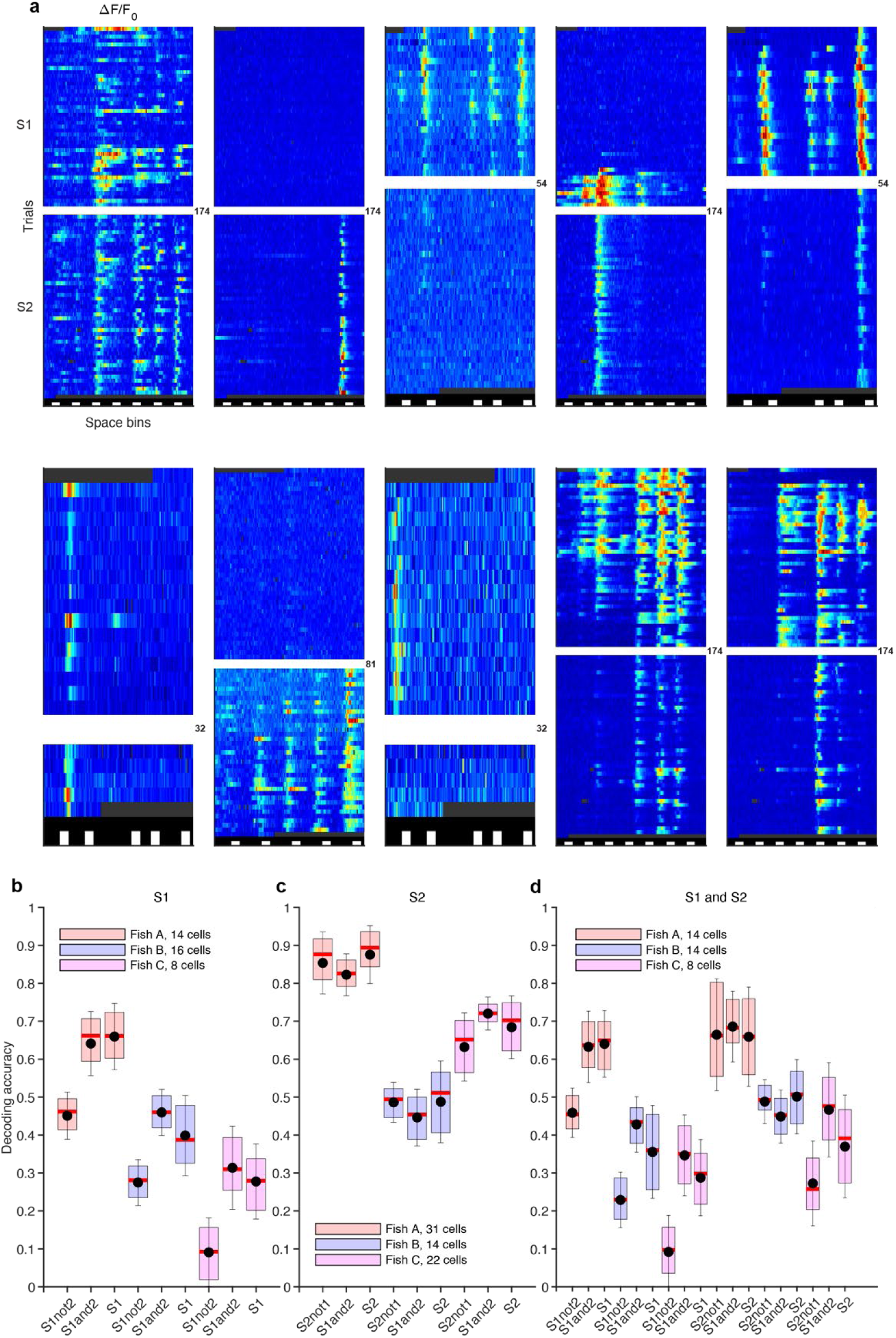
Evolution of the internal representation from S1 to S2. **a,** Additional examples of spatial activity maps of individual neurons in S1 and S2. Conventions as in Fig. 1h. Sessions are separated by white spacing, numbers indicate trials between sessions. **b** - **d,** Decoding accuracy using neuronal subsets of different size for each of three fish imaged in S1 and S2. Subset sizes are indicated and given by the smallest number of spatially selective neurons in any of the categories in each fish. Dots and error bars show mean and s.d., respectively; box plots show median and surrounding quartiles.

**Extended Data Fig. 4.**
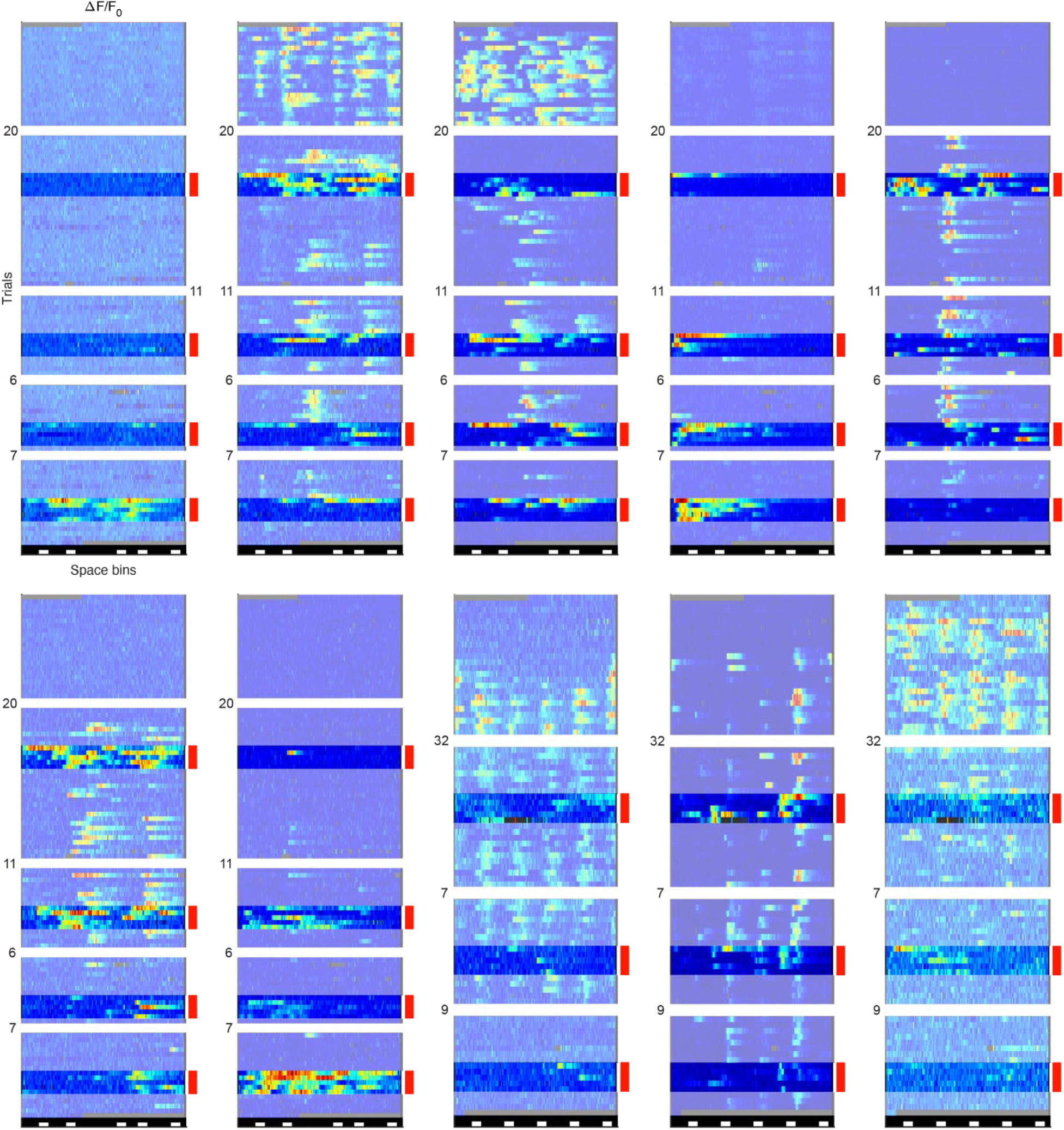
Additional examples of effects of landmark deletion on neuronal activity. Additional examples of spatial activity maps of different neurons across all imaging sessions in experiments including landmark deletions. Sessions are separated by white spacing, numbers indicate trials between sessions. Red bars and stronger colors highlight trials when landmarks were deleted. Conventions as in Fig. 1h.

**Extended Data Fig. 5.**
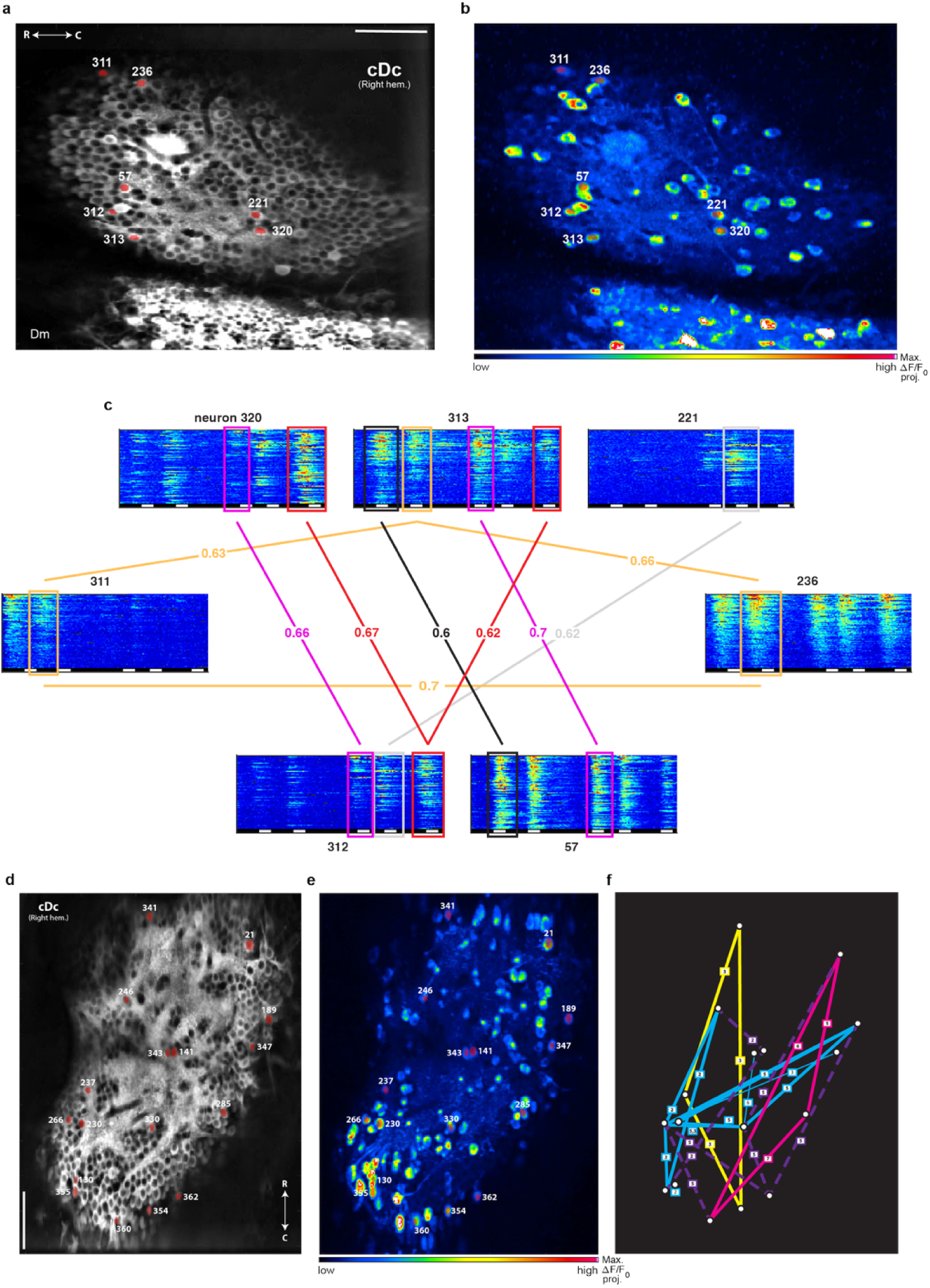
Additional examples of functional neuronal ensembles in cDc. **a,** Raw fluorescence image (time-averaged) of neurons in cDc with regions of interest (ROIs) corresponding to neurons of identified ensembles in a fish in the 5Irr environment. Right hem., right hemisphere; R, rostral; C, caudal; numbers show neuron IDs; scale bar, 50 μm. **b,** Maximum projection of **Δ**F/F_0_ image time-series. Annotations show neurons of identified ensembles. **c,** Spatial activity maps of ensemble neurons highlighted in **a** and **b**. Colored boxes and lines highlight inter-neuron field pairs with high positive noise correlations. Correlation coefficients are indicated. Colors correspond to different VR sectors. **d**, **e**, Neurons associated with multiple ensembles in another fish in the 7Reg environment, overlaid on raw fluorescence image and maximum **Δ**F/F_0_ projection. **f**, Schematic of identified functional neuronal ensembles. Solid lines show high positive correlations within three neuronal ensembles (yellow, blue, and magenta) between neurons (white dots); dashed purple lines show high negative correlations. Boxed numbers indicate VR sectors associated with the corresponding fields.

